# Inhibitory tagging both speeds and strengthens saccade target selection in the superior colliculus during visual search

**DOI:** 10.1101/2023.09.20.558470

**Authors:** Christopher Conroy, Rakesh Nanjappa, Robert M. McPeek

## Abstract

It has been suggested that, during difficult visual search tasks involving time pressure and multiple saccades, inhibitory tagging helps to facilitate efficient saccade target selection by reducing responses to objects in the scene once they have been searched and rejected. The superior colliculus (SC) is a midbrain structure involved in target selection, and recent findings suggest an influence of inhibitory tagging on SC activity. Precisely how, and by how much, inhibitory tagging influences target selection by SC neurons, however, is unclear. The purpose of this study, therefore, was to characterize and quantify the influence of inhibitory tagging on target selection in the SC. Rhesus monkeys performed a visual search task involving time pressure and multiple saccades. Early in the fixation period between saccades, a subset of SC neurons reliably discriminated the stimulus selected as the next saccade goal, consistent with a role in target selection. Discrimination occurred earlier and was more robust, however, when unselected stimuli in the search array had been previously fixated on the same trial. This indicates that inhibitory tagging both speeds and strengthens saccade target selection in the SC during multisaccade search. The results provide constraints on models of target selection based on SC activity.

**NEW AND NOTEWORTHY:** An important aspect of efficient behavior during difficult, time-limited visual search tasks is the efficient selection of sequential saccade targets. Inhibitory tagging, i.e., a reduction of neural activity associated with previously fixated objects, may help to facilitate such efficient selection by modulating the selection process in the superior colliculus (SC). In this study, we characterized and quantified this modulation and found that, indeed, inhibitory tagging both speeds and strengthens target selection in the SC.

## INTRODUCTION

Our daily lives are replete with examples of visual search tasks that we need to complete efficiently. Under time pressure, we execute saccade sequences, interleaved with brief fixations, to search the visual scene for objects of interest. Performance in such contexts depends critically on efficient and appropriate selection of sequential saccade targets. Inhibitory tagging, i.e., a reduction of neural activity associated with previously fixated objects, may help to facilitate such efficient selection by increasing the speed and strength with which uninspected objects can be discriminated from previously fixated objects in the visual scene (1–6).

In a recent study, we reported evidence of an influence of inhibitory tagging on activity in the superior colliculus (SC) (6), a midbrain structure that is involved in saccade target selection (7–25). Specifically, we found that, during a time-limited, multisaccade visual search task, the activity of many SC neurons was lower when the stimuli in their response fields (RFs) had been previously fixated compared to when they had not. Moreover, the largest effects of inhibitory tagging were observed in the subset of SC neurons that had response properties consistent with a role in saccade target selection. This suggests that modulation of the selection process at the level of individual SC neurons may provide a mechanism by which inhibitory tagging supports efficient search behavior. Precisely how, and by how much, inhibitory tagging influences target selection in the SC, however, is unclear, as we did not examine nor quantify the influence of inhibitory tagging on target selection in our previous study.

The purpose of this study, therefore, was to extend our previous work and to characterize and quantify the influence of inhibitory tagging on target selection in the SC. As an approach, we analyzed the ability of SC neurons to discriminate between selected and unselected RF stimuli during the brief fixation period between saccades in the multisaccade search task and compared the time course and strength of discrimination with and without inhibitory tagging. More specifically, we first analyzed the ability of SC neurons to discriminate between RF stimuli that were selected as the saccade target and unselected stimuli in the absence of inhibitory tagging, i.e., when neither the selected nor unselected stimuli had been previously fixated on the same trial. We then compared the time course and strength of discrimination in this condition with results obtained when the unselected stimuli had been tagged with inhibition due to previous fixation on the same trial. We reasoned that if inhibitory tagging supports efficient search behavior by modulating the selection process in the SC, we should observe more rapid and robust discrimination when the unselected stimuli had been tagged with inhibition. As will be seen, this is what we found, indicating that inhibitory tagging both speeds and strengthens saccade target selection in the SC.

## METHODS

Experimental protocols were approved by the Institutional Animal Care and Use Committee at SUNY College of Optometry and complied with the guidelines of the US Public Health Service policy on Humane Care and Use of Laboratory Animals. A detailed description of the surgical, recording, and behavioral procedures can be found in our recent report (6). Briefly, we used microelectrodes and standard methods to record the activity of well-isolated single SC neurons in two male rhesus monkeys (*Macaca mulatta*). During recording, monkeys performed two behavioral tasks for liquid reward: a delayed-saccade task and a multisaccade search task. Eye movements were measured using a video-based eye tracker (Eyelink 1000, SR Research).

The delayed-saccade task was used to determine the RF center for each neuron and for neuronal classification. At the beginning of each trial of this task, a red or green disc appeared at the center of a stimulus-display monitor. Monkeys were required to fixate this central disc for 450-650 ms, at which point a red or green target disc appeared at a peripheral location. Following the appearance of the target disc, monkeys were required to maintain fixation for an additional 500-700 ms, at which point the central disc disappeared. Monkeys were rewarded for making a saccade to the target disc within 70-400 ms of the central disc’s disappearance. All discs subtended 0.5-2.0° of visual angle [M-scaled based on eccentricity to keep salience approximately constant (cf. Ref. 26)] and had a luminance of 12 cd/m^2^ on a 0.3 cd/m^2^ background. The color of the target disc always differed from that of the central disc, was randomly selected at the beginning of each recording session and was fixed throughout each session.

Initially, the location of the target was manually varied to home in the neuron’s RF center, defined as the location yielding the largest visual and/or motor response. Following identification of the RF center, the activity of each neuron was recorded in at least ten trials of the delayed-saccade task with the target positioned at the RF center.

At the beginning of each trial of the search task, a red or green disc appeared at the center of the monitor. Monkeys were required to fixate this disc for 500 ms, at which point a search array appeared. The array consisted of multiple red and green discs positioned at the intersection points of an invisible grid centered on the central disc. All discs had the same size and luminance as the discs presented to the RF center in the delayed-saccade task. One of the discs in the array (never the central disc) held a reward, meaning that, when fixated, a liquid reward was immediately administered to the monkey. The color of the reward-bearing disc always differed from the color of the central disc and, within a session, remained constant from trial to trial. Thus, if the central disc was green, the reward-bearing disc was red, and vice versa. No other colors were tested. Each trial lasted for eight seconds or until the reward-bearing disc had been fixated, whichever came first. For each neuron, the array was configured such that when the monkey fixated one of the discs, a nearby disc was positioned at the RF center. For most neurons, this positioning scheme permitted the use of a five-by-five rectangular array and, for most of these neurons, the array consisted of nine potentially reward bearing discs. For some neurons, typically with large RF eccentricities, the array consisted of fewer discs, in which cases attempts were made to maintain approximately the same ratio of potentially reward-bearing to not-potentially reward-bearing discs as in the five-by-five rectangular array case (e.g., for one neuron with an RF center at 15°, the array consisted of 17 discs, six of which were potentially reward bearing). The locations of potentially reward-bearing and not-potentially reward-bearing discs within the array were randomly selected on each trial. On a small proportion of all trials (roughly 7%), all “potentially” reward-bearing discs were in fact reward bearing, meaning that the monkey received a reward upon the first (and only the first) fixation of each disc that had a color that differed from that of the central disc. This trial type was used to increase the monkey’s motivation. These trials, like the single-reward trials, lasted for eight seconds or until all reward-bearing discs had been fixated, whichever came first.

### Data analysis

As noted in the Introduction, the purpose of this study was to examine the influence of inhibitory tagging on target selection in the SC. Thus, we focused our analyses on neurons exhibiting significant delay-period activity in the delayed-saccade task because SC neurons involved in saccade target selection tend to exhibit this type of activity (cf. 7-12, 18, 21, 24, 25). To measure delay-period activity, we compared each neuron’s activity during a 100-ms window beginning 300-ms after the onset of the target to its baseline activity during a 100-ms window beginning 100-ms before target onset. Only trials in which the target was positioned at the RF center were considered. If delay-period activity was significantly greater than baseline activity (Wilcoxon rank-sum test, *α* = 0.05), the neuron was included in subsequent analyses.

For the search task, the activity of each neuron was analyzed during fixations between sequential saccades. Activity was analyzed separately for three categories of fixation, which differed in terms of the fixation status of the RF disc [previously fixated (PF) or not previously fixated (NPF) in the same trial] and whether the RF disc was selected as the next saccade goal: (1) fixations during which an NPF disc in the RF was selected as the next saccade goal (NPF-in-RF, saccade-towards); (2) fixations during which an NPF disc in the RF was not selected as the next saccade goal (NPF-in-RF, saccade-away); and (3) fixations during which a PF disc in the RF was not selected as the next saccade goal (PF-in-RF, saccade-away). A neuron was only included in the analyses if its activity was recorded during more than five fixations for each category.

The initial fixation in each trial did not count towards this criterion and, moreover, was not included in any analysis. Schematic illustrations of the search task, the search array, an example saccade sequence, and the three fixation categories are shown in Fig. 1.

**FIG. 1:**
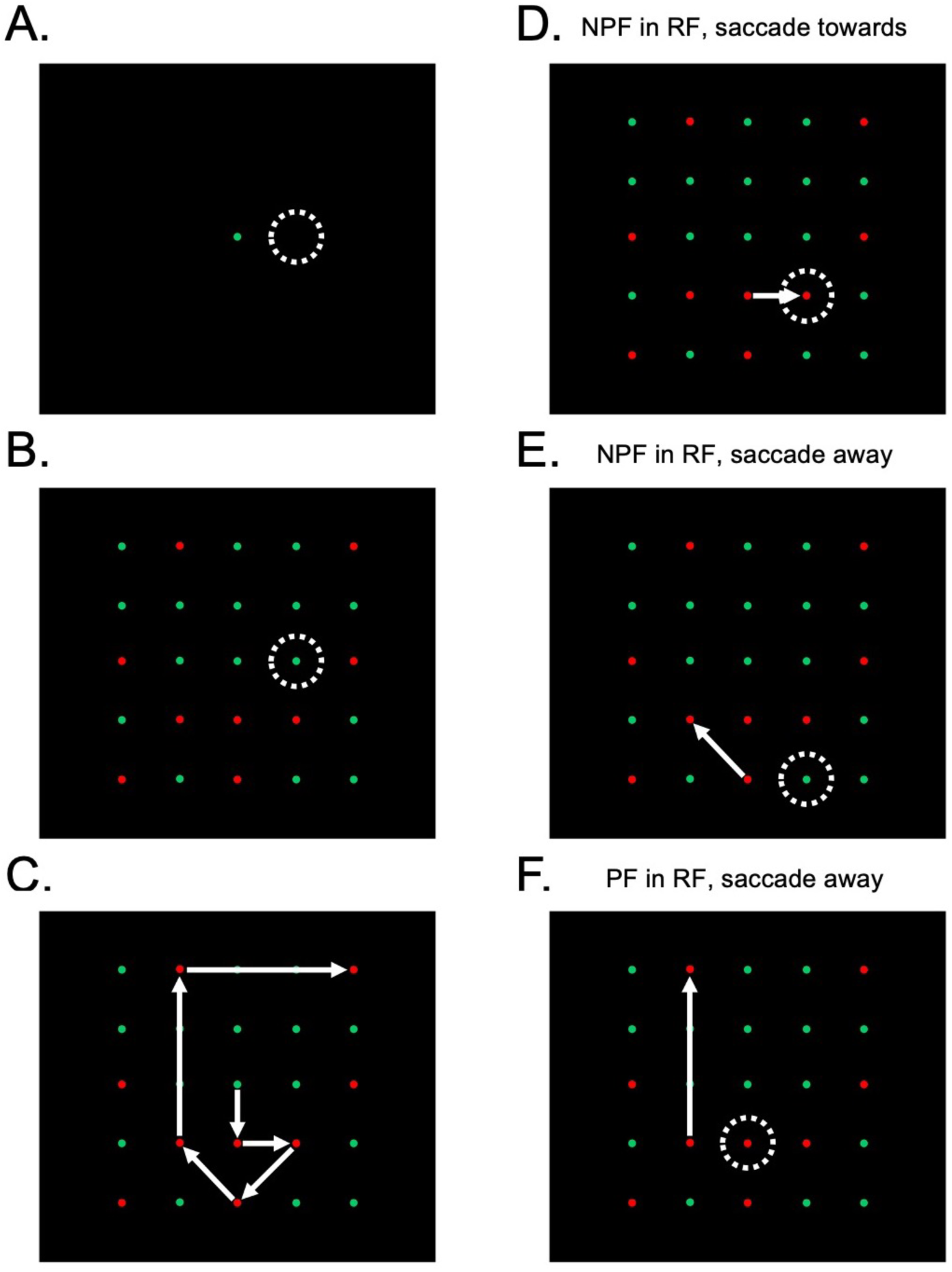
Schematic illustrations of the search task. **A.** Trials began with a disc at display center which monkeys fixated. **B.** The search array then appeared on the screen. At fixation, a nearby disc was positioned at the center of the RF of the neuron under study (dotted white circle). One disc differing in color from the central disc triggered a liquid reward when fixated. The monkey’s task was to fixate this reward-bearing disc. **C.** Fixation of the reward-bearing disc was typically preceded by a sequence of multiple saccades to other discs (see Ref. 6 for details on search behavior). This panel illustrates a situation where fixation of the reward-bearing disc (in this example, the disc in the upper right corner) was preceded by a sequence of six saccades, represented by white arrows. Analyses focused on three fixation categories: **D.** fixations during which an NPF disc in the RF was selected as the next saccade goal (NPF-in-RF, saccade-towards), **E.** fixations during which an NPF disc in the RF was not selected as the next saccade goal (NPF-in-RF, saccade-away), and **F.** fixations during which a PF disc in the RF was not selected as the next saccade goal (PF-in-RF, saccade-away). In **D-F**, the arrow illustrates the vector of the next saccade and the dotted-white circle illustrates the RF center.

We recorded 113 neurons from two monkeys. Of these, 65 met our inclusion criteria. For each neuron, spike-density functions were created by convolving individual spikes with a Gaussian kernel (*σ* = 10 ms). We then applied two receiver operating characteristic (ROC) analyses (27) to the activity of each neuron as a way to measure the effects of inhibitory tagging on the target-selection process. Similar ROC analyses have been used in previous studies of target selection in the SC and other areas (e.g., Refs. 9-12, 16, 18, 19, 21, 24, 25, 28-30). The first analysis measured the discriminability of selected and unselected RF discs in the absence of inhibitory tagging by comparing activity during NPF-in-RF, saccade-towards fixations to the activity during NPF-in-RF, saccade-away fixations. We refer to this as the NPF-re-NPF analysis. The other analysis measured how inhibitory tagging of unselected discs affects target selection in the SC by comparing activity during NPF-in-RF, saccade-towards fixations to activity during PF-in-RF, saccade-away fixations. We refer to this as the NPF-re-PF analysis. We used the same NPF-in-RF, saccade-towards activity for both analyses because, as noted in the Introduction, we were interested in understanding how inhibitory tagging supports efficient search behavior by modulating SC target selection. In this task, efficient search requires selection of NPF discs. Thus, if inhibitory tagging supports efficient search behavior by modulating the selection process in the SC, it should facilitate efficient selection of NPF discs in particular.

For both ROC analyses, activity was aligned on both fixation onset and the onset of the subsequent saccade (fixation offset). For each alignment, an ROC curve was computed at each ms by comparing the distributions of activity for selected and unselected disc, and area under the ROC curve was computed. An ROC area of 0.5 indicates complete overlap between selected and unselected distributions, areas closer to 1.0 indicate greater activity for selected discs, and areas closer to 0.0 indicate greater activity for unselected discs. We analyzed both fixation- and saccade-onset-aligned activity to accurately measure the time course of target discrimination relative to both fixation and saccade onset, as has been standard in previous studies (e.g., 9-11, 16, 18, 24, 25, 28-30). Statistical significance was assessed using a permutation test (2,000 iterations) on ROC area (cf. 9, 10, 16, 21, 30) at each ms beginning 50 ms before and ending 150 ms after fixation onset for the fixation-onset-aligned analysis and beginning 150 ms before and ending with saccade onset for the saccade-onset-aligned analysis.

For all ROC analyses, we obtained two selection metrics for each neuron: discrimination time and magnitude (e.g., 18, 24, 25, 29). For the fixation-onset-aligned analysis, discrimination time was defined as the earliest time point within the analysis window at which ROC area was significantly different from 0.5, the mean selected activity was greater than the mean unselected activity, and both of these conditions were met for at least 20 consecutive ms. For the saccade-onset alignment, discrimination time was defined as the earliest time within the analysis window at which ROC area was significantly different from 0.5, the mean selected activity was greater than the mean unselected activity, and both of these conditions were met at all time points until saccade onset. Discrimination magnitude was defined as the maximum ROC area within each analysis window. Thus, the discrimination-time metric provided an estimate of the earliest time point at which each neuron reliably discriminated selected from unselected discs in its RF and therefore an estimate of the time of the completion of the selection process. We predicted that if inhibitory tagging supports efficient search behavior by modulating the selection process in the SC, discrimination of an NPF target disc should be faster when that target is embedded in PF discs (NPF-re-PF analysis) than when it is embedded in NPF discs (NPF-re-NPF analysis). In other words, we anticipated shorter discrimination times for the NPF-re-PF than NPF-re-NPF analysis. Discrimination magnitude provided an estimate of the discriminability of selected and unselected discs following completion of the selection process. We predicted that if inhibitory tagging strengthens selection signals in the SC, it should increase discriminability of selected and unselected discs when the unselected discs have been previously fixated. Thus, we anticipated larger discrimination magnitudes for the NPF-re-PF than NPF-re-NPF analysis. Statistical comparisons of selection metrics were accomplished using Wilcoxon signed-rank tests (*α* = 0.05).

## RESULTS

Many of the neurons in our sample exhibited activity during the search task consistent with a role in saccade target selection, and, for many of these neurons, the time course of the selection process, as well as the strength of selection signals, was influenced by inhibitory tagging. This is illustrated in Fig. 2, which shows the results (activity and ROC area versus time) for an example neuron. We focus first on the activity associated with NPF RF discs and the results of the NPF-re-NPF analysis, which, again, characterized the selection process without inhibitory tagging. Relatively early in the fixation period, this neuron reliably discriminated selected from unselected NPF discs. The fixation- and saccade-onset-aligned discrimination times for the NPF-re-NPF analysis were 53 ms relative to fixation onset and -112 ms relative to saccade onset, respectively. Activity associated with selected discs steadily increased and eventually culminated in a saccadic burst time locked to the next saccade, whereas activity associated with unselected discs was initially inhibited and then held at bay. The consequence of these two activity patterns was robust selected-unselected discriminability (i.e., NPF-re-NPF ROC areas > 0.5) throughout most of the fixation period following the initial phase of discrimination and, in particular, around the time of saccade onset. Discrimination magnitudes for the NPF-re-NPF analysis aligned on fixation and saccade onset, which, it can be seen, reflected ROC areas around the time of saccade onset, were 0.76 and 0.83, respectively. Thus, before inhibitory tagging, this neuron reliably and robustly discriminated selected from unselected NPF discs, consistent with a role in saccade target selection.

**FIG. 2:**
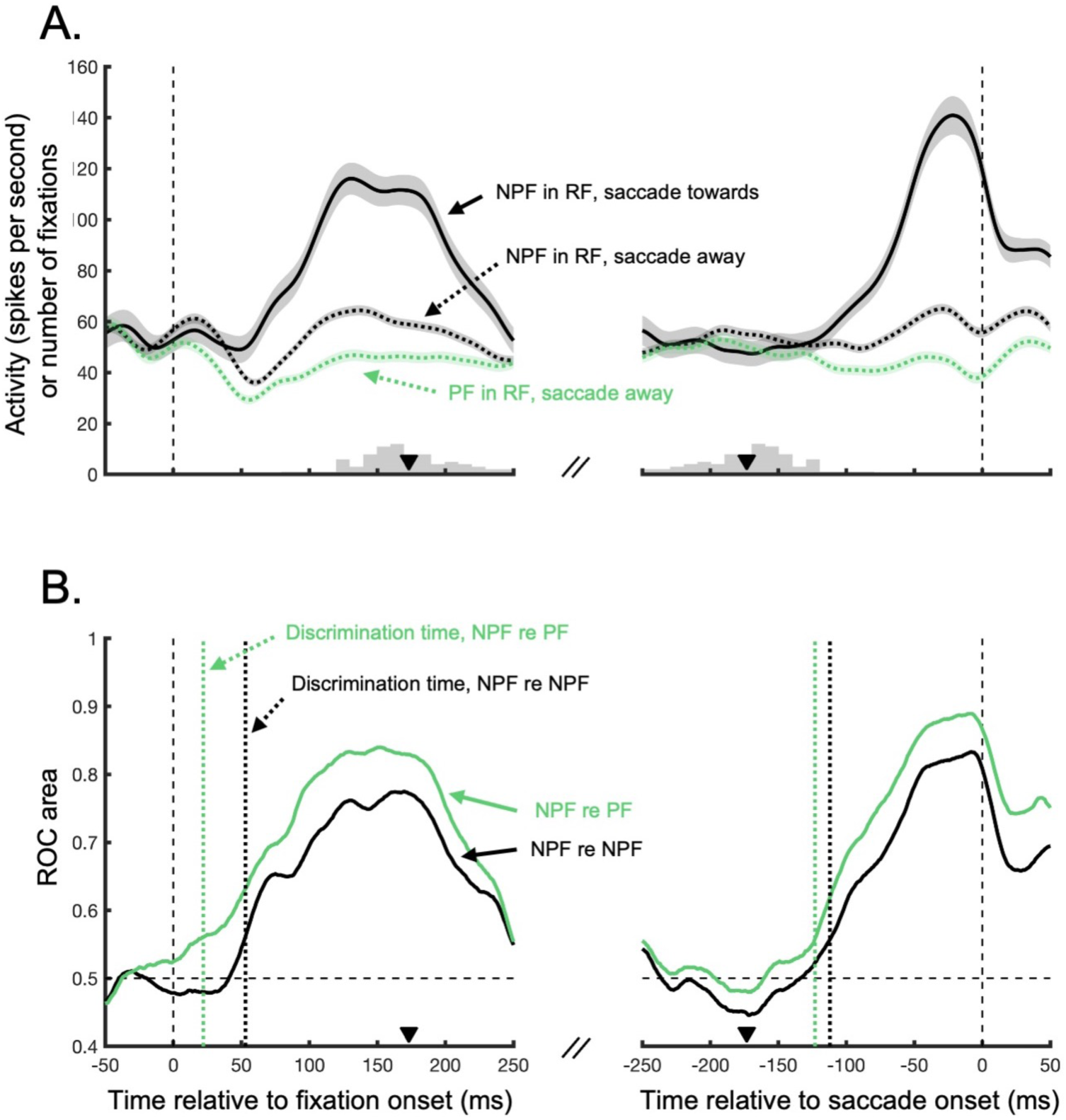
Activity and ROC area functions for an example neuron. **A.** Mean fixation- (left panel) and saccade-onset- (right panel) -aligned spike-density functions. Different traces show activity for different fixation categories: the solid black trace shows the activity during NPF-in-RF, saccade-towards fixations, the dotted black trace shows the activity during NPF-in-RF, saccade-away fixations, and the dotted green trace shows the activity during PF-in-RF, saccade-away fixations. Each trace represents the mean activity and the shaded region represents +/-1 standard error of the mean. In the left column, the inset histogram shows the distribution of saccade-onset times for the NPF-in-RF saccade-towards fixations. In the right column, it shows the distribution of fixation onset times for the same fixation category. The y-axis in each panel gives the fixation counts for the distributions [i.e., the numbers on the y axis give both spikes per second (for the spike-density functions) and fixation counts (for the fixation-onset time distribution)]. The downward pointing triangles show the means of the distributions. A small number of fixations were longer than 250 ms and are not shown for visualization purposes. **B.** Fixation- (left panel) and saccade-onset- (right panel) -aligned ROC area functions for the same neuron. The black trace shows the results of the NPF-re-NPF analysis, the green trace shows the results of the NPF-re-PF analysis, and the vertical dotted lines show the discrimination times for the two analyses. The downward pointing triangles replot the mean saccade onset (left column) and onset (right column) times for NPF-in-RF, saccade-towards fixations from Fig. 2A.

Discrimination of selected NPF discs occurred earlier and was more robust, however, following the placement of inhibitory tags on unselected discs in the search array, as indicated by the decrease in discrimination times and increase in discrimination magnitudes for the NPF-re-PF relative to the NPF-re-NPF analysis. For the fixation-onset-aligned analysis, the discrimination time decreased from 53 ms for the NPF-re-NPF condition to 22 ms for the NPF-re-PF condition, i.e., when the unselected stimulus had been tagged with inhibition. Furthermore the discrimination magnitude increased from 0.76 to 0.84 going from the NPF-re-NPF to NPF-re-PF analysis. A decrease in discrimination time (from -112 ms to -123 ms for the NPF-re-NPF to NF-re-PF analysis) and increase in discrimination magnitude (from 0.83 to 0.89 for the NPF-re-NPF to NF-re-PF analysis) occurred for the saccade-onset-aligned analysis as well. The discrimination magnitudes for the NPF-re-PF analysis also reflected ROC areas around the time of saccade onset, and so the larger discrimination magnitude for the NPF-re-PF than NPF-re-NPF analysis indicates that inhibitory tagging of unselected discs was strong even around the time of saccade onset, potentially reducing the probability that activity associated with unselected discs would disrupt or interfere with the selected saccade (31, 32).

A decrease in discrimination times and increase in discrimination magnitudes after inhibitory tagging was evident for many of the neurons in our sample. Figure 3 shows the fixation-onset-aligned target-selection metrics for the 51 of 65 neurons in our sample that discriminated selected from unselected RF discs during the fixation-onset-aligned analysis window (i.e., neurons that yielded a fixation-onset-aligned discrimination time for both the NPF-re-NPF and NPF-re-PF analyses). For these neurons and alignment, the median discrimination time was significantly shorter for the NPF-re-PF than NPF-re-NPF analysis (33 versus 49 ms relative to fixation onset; *Z* = - 3.953; *P* < .001; Fig. 3A-B). Indeed, for the NPF-re-PF analysis, many neurons exhibited very early discrimination times, with a substantial number discriminating selected from unselected discs even before fixation onset and/or the visual-afferent delay. Specifically, for the NPF-re-PF fixation-onset-aligned analysis, 19 of the 51 neurons that discriminated selected from unselected discs did so before fixation onset. An additional 15 did so within the first 50 ms following fixation onset. Such early discrimination times are consistent with the idea that the discrimination process began before fixation onset (i.e., during the planning or execution of the previous saccade; cf. Refs. 25, 33) and that there was a predictive remapping of inhibitory-tagging signals preceding the onset of each saccade (cf. Refs 5, 6). Thus, inhibitory tagging may not only facilitate the selection process during individual fixations in multisaccade sequences but also may facilitate the efficient planning of multisaccade sequences as a whole.

**FIG. 3:**
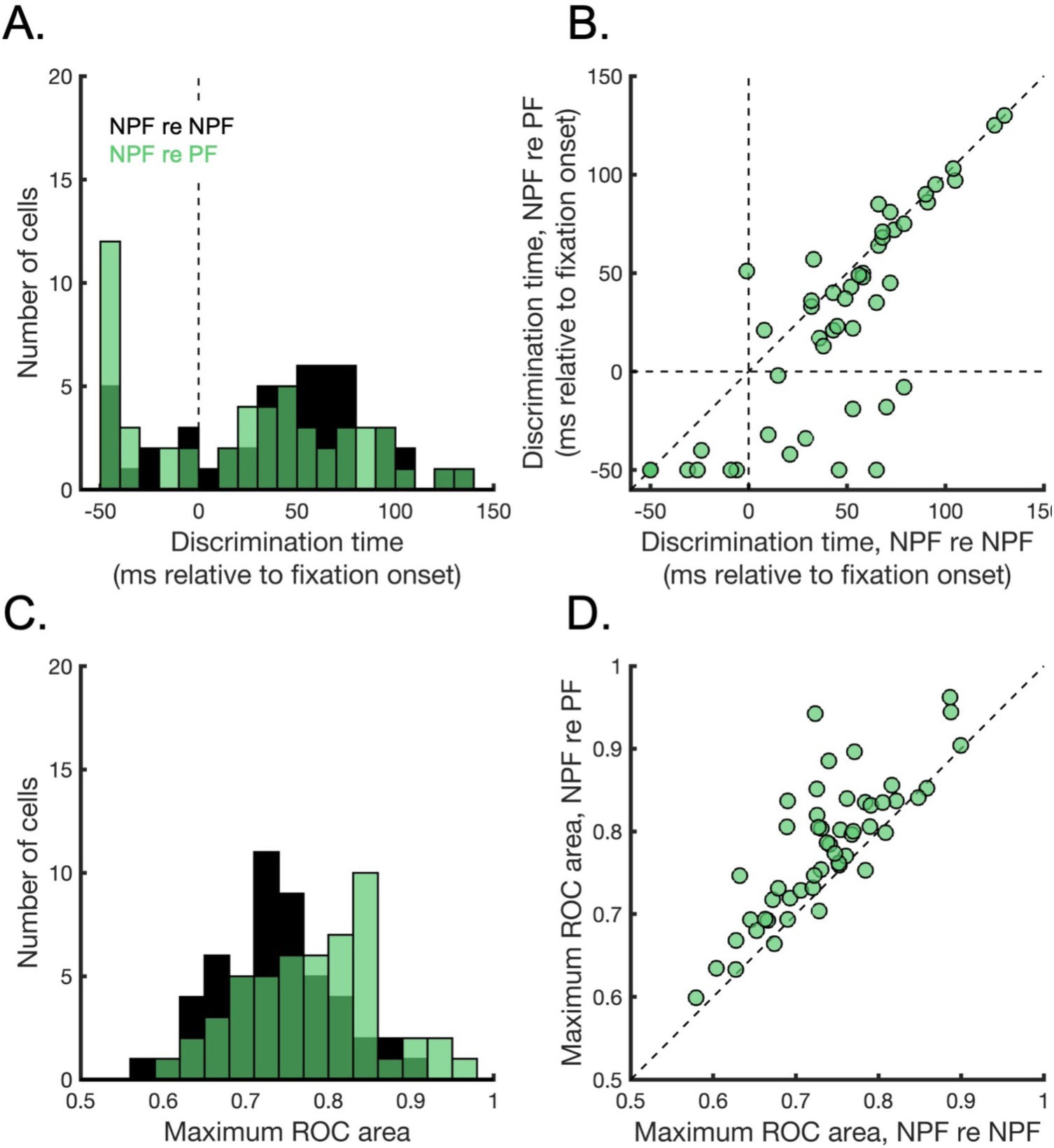
Summary of the fixation-onset-aligned target-selection metrics for the 51 of 65 neurons in our sample that discriminated the saccade target within the fixation-onset-aligned analysis window for both the NPF-re-NPF and NPF-re-PF analyses. **A.** Distributions of discrimination times for the two analyses. The black and green bars show the distributions for the NPF-re-NPF and NPF-re-PF analyses, respectively. Note that -50 ms relative to fixation onset was the earliest possible discrimination time given the analysis window. **B.** Relationship between NPF-re-PF and NPF-re-NPF discrimination times for individual neurons (individual symbols). **C.** Distributions of discrimination magnitudes for the NPF-re-NPF (black bars) and NPF-re-PF (green bars) analyses. **D.** Relationship between NPF-re-PF and NPF-re-NPF discrimination magnitudes for individual neurons (individual symbols).

For the same subset of 51 neurons that discriminated selected from unselected discs during the fixation-onset-aligned analysis window, the median discrimination magnitude was higher for the NPF-re-PF than NPF-re-NPF analysis (0.79 versus 0.73; *Z* = 5.521; *P* < .001; Fig. 3C-D). For the 48 neurons that discriminated selected from unselected RF discs during the saccade-onset-aligned analysis window, we observed a similar pattern: the median discrimination time was shorter (-113 versus -97 ms relative to saccade onset; *Z* = -2.962; *P* = .003; Fig. 4A-B) and the median discrimination magnitude was larger (0.83 versus 0.79; *Z* = 4.636 *P* < .001; Fig. 4C-D) for the NPF-re-PF than NPF-re-NPF analysis. Thus, we conclude that inhibitory tagging both speeds and strengthens saccade target selection at the level of individual SC neurons.

**FIG. 4:**
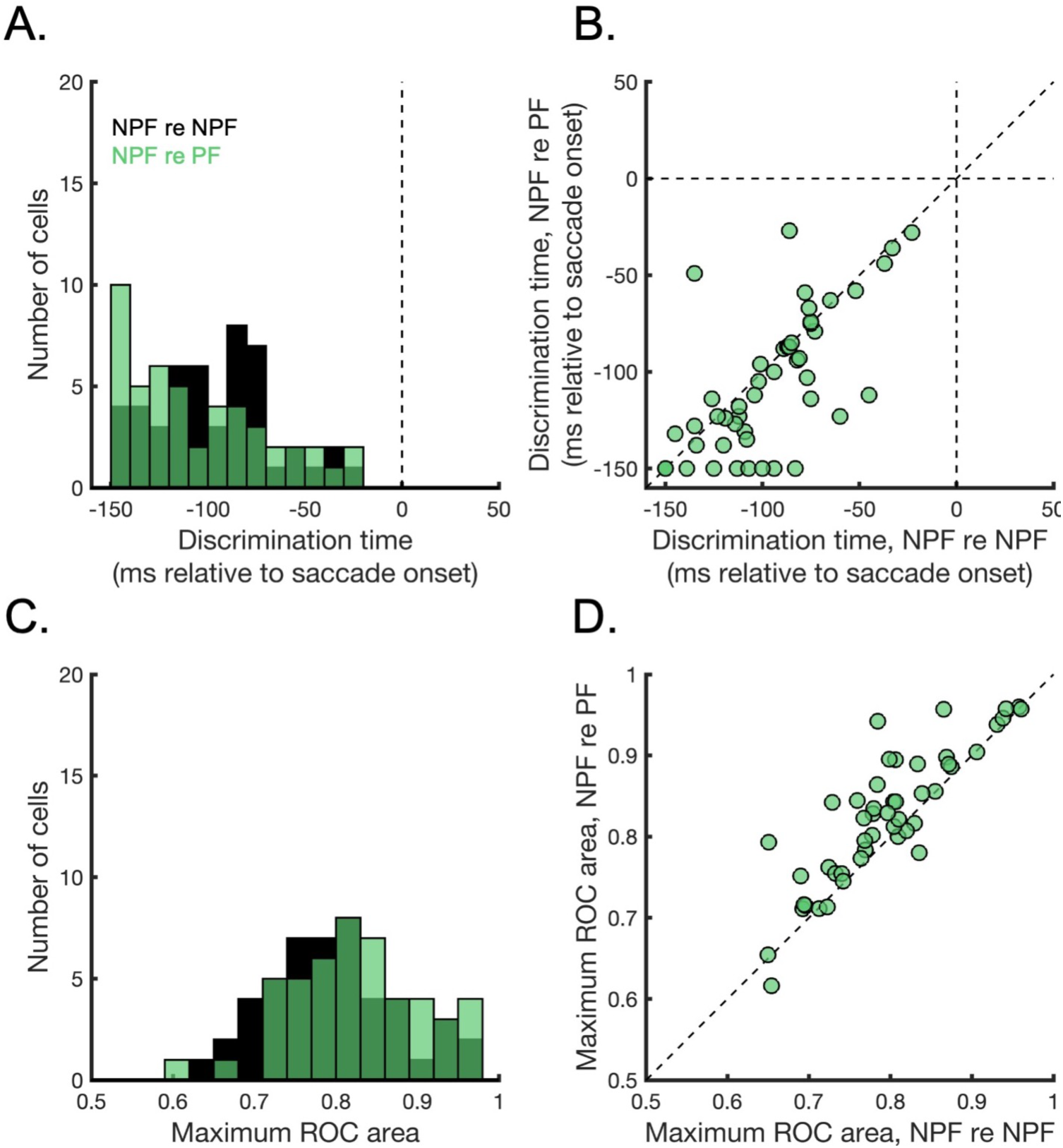
Summary of the saccade-onset-aligned target-selection metrics for the 48 of 65 neurons in our sample that discriminated the saccade target within the saccade-onset-aligned analysis window (-150-0 ms relative to saccade onset) for both the NPF-re-NPF and NPF-re-PF analyses. Otherwise, what is shown in each panel corresponds to what is shown in Fig. 3 and the plotting conventions are the same. Note that, for the saccade-onset-aligned discrimination times, however, the earliest possible discrimination time was -150 ms.

Additional analyses that bolster this conclusion are reported in the supplementary material. Also in the supplementary material, we report analyses showing a relationship between behavioral search efficiency during the SC recording sessions and the improvement in selection metrics for the SC neurons recorded during those sessions. This supports the view that the target-selection benefits shown here contributed to efficient search behavior.

## DISCUSSION

Saccade target selection involves a competition between potential saccade goals on the SC visuomotor map. An inhibitory-tagging mechanism that facilitates efficient visual search would be expected to influence this competitive process by reducing activity associated with previously fixated objects and thus biasing the competitive process towards uninspected objects in the visual scene. Previously, we showed this basic pattern of modulations after inhibitory tagging in the SC (6), perhaps reflecting the influence of inhibitory-tagging signals originating in cortical areas (4, 5). Here, we characterized and quantified the influence of these modulations on SC target selection. Using ROC analyses, we found that inhibitory tagging both speeds and strengthens saccade target selection in the SC, yielding significantly faster discrimination of selected and unselected RF stimuli and significantly larger discrimination magnitudes. We also found that the benefit of inhibitory tagging for target selection in these neurons was related to the monkeys’ behavioral search efficiency during the SC recording sessions.

Thus, we conclude that modulation of the selection process at the level of individual SC neurons likely provides a mechanism by which inhibitory tagging supports efficient search behavior. Models of target selection based on SC (or, more generally, priority-map) activity (e.g., Ref. 34) may wish to take these modulations into account to more accurately capture the dynamics of the selection process during time-limited, multisaccade search.

## SUPPLEMENTARY MATERIAL

Supplementary material: DOI: https://doi.org/10.6084/m9.figshare.24596952

## Acknowledgements

This work was funded by National Institutes of Health (NIH)- National Eye Institute (NEI) grant number R01-EY030669

