## Supplementary material for "Inhibitory tagging both speeds and strengthens saccade target selection in the superior colliculus during visual search"

### *Influence of RF disc color on SC target-selection metrics*

To determine if the color of the RF disc—meaning, its color in the context of the task: potentially reward bearing or not—had an influence on the results, we performed the same ROC analyses as reported in the main article on two restricted datasets: one consisting of fixations during which the RF disc was always potentially reward bearing and another consisting of fixations during which the RF disc was always not potentially reward bearing. In both cases, discrimination times decreased and discrimination magnitudes increased following inhibitory tagging.

More specifically, for the dataset limited to fixations during which the RF disc was potentially reward bearing, there was a significant decrease in discrimination times and increase in discrimination magnitudes after inhibitory tagging for both the fixation- and saccade-onset-aligned analyses. For the 45 neurons from this dataset that discriminated selected from unselected RF discs during the fixation-onset-aligned analysis window, discrimination times decreased from 62 to 27 ms relative to fixation onset ($Z = -4.632$; $P < .001$), and discrimination magnitudes increased from 0.74 to 0.80 ($Z = 4.984$; $P < .001$), from the NPF-re-NPF to the NPF-re-PF analysis. For the 41 neurons that discriminated selected from unselected discs during the saccade-onset-aligned analysis window, discrimination times decreased from -70 ms to -97 ms relative to saccade onset ($Z = -3.630$; $P < .001$) and discrimination magnitudes increased from 0.81 to 0.84 ($Z = 4.970$; $P < .001$).

Similarly, for the dataset limited to fixations during which the RF disc was not potentially reward bearing, inhibitory tagging yielded a decrease in discrimination times and increase in discrimination magnitudes. For the 35 neurons from this dataset that discriminated selected from unselected RF discs during the fixation-onset-aligned analysis window, discrimination times decreased from 59 to 45 ms from the NPF-re-NPF to the NPF-re-PF analysis ($Z = -2.245$; $P = .025$) and discrimination magnitudes increased from 0.75 to 0.81 ($Z = 4.128$; $P < .001$). For the 35 neurons that discriminated selected from unselected RF discs during the saccade-onset-aligned analysis window, there was a significant increase in discrimination magnitudes (from 0.80 to 0.85 from the NPF-re-NPF to NPF-re-PF analysis; $Z = 3.259$; $P = .001$) and clear trend towards earlier discrimination times (from -74 to -91 relative to saccade onset from the NPF-re-NPF relative to NPF-re-PF analysis; $Z = -1.936$; $P = .053$).

Thus, we conclude that the color of the RF disc did not affect the overall pattern of results or our conclusions regarding the influence of inhibitory tagging on target selection in the SC.

### *Influence of inhibitory tagging on SC motor bursts*

To determine if inhibitory tagging had an influence on SC motor bursts for saccades made to PF versus NPF targets in the RF, we compared activity during NPF-in-RF, saccade-towards fixations to activity during PF-in-RF, saccade-towards fixations (i.e., fixations during which the RF disc had been previously fixated on the same trial and was selected as next saccade goal, a category of fixation not analyzed in the main article).

Figure S1A shows the mean activity across the 65 neurons in our sample for these two categories of fixation aligned on fixation onset (Fig. S1A, left panel) and saccade onset (Fig. S1A, right panel). During the early visual period immediately following fixation onset, there was a slight difference in activity between these two categories of fixation, with activity during PF-in-RF, saccade-towards fixations being slightly lower than activity during NPF-in-RF, saccade-towards fixations (Fig. S1A, left panel). This is consistent with an influence of inhibitory tagging on visually evoked activity regardless of the ultimate choice of saccade goal. Around the time of saccade onset, activity during the two categories of fixation was indistinguishable (Fig. S1A, right panel), suggesting that inhibitory tagging does not affect SC motor bursts for saccades made into the RF.

These impressions are confirmed by the results of an ROC analysis, which compared NPF-in-RF, saccade-towards activity to PF-in-RF, saccade towards activity (Fig. S1B). On average, ROC areas were slightly above 0.5 early in the fixation period (Fig. S1B, left panel), but decreased and hovered around 0.5 by the time of the motor burst (Fig. S1B, right panel). Indeed, for the fixation-onset-aligned ROC analysis comparing NPF-in-RF to PF-in-RF saccade-towards activity, 12 of the 65 neurons yielded a discrimination time (see main article for statistical criterion). For these 12 neurons, discrimination times ranged from -50 to 108 ms relative to fixation onset and discrimination magnitudes ranged from 0.59 to 0.84. For the saccade-onset-aligned ROC analysis, however, none of the 65 neurons yielded a discrimination time, indicating that they did not exhibit significantly different activity throughout the motor burst during NPF-in-RF and PF-in-RF saccade-towards fixations.

Thus, we conclude that inhibitory tagging primarily affects visually evoked activity in the SC and the target-selection process via an inhibition of unselected activity.

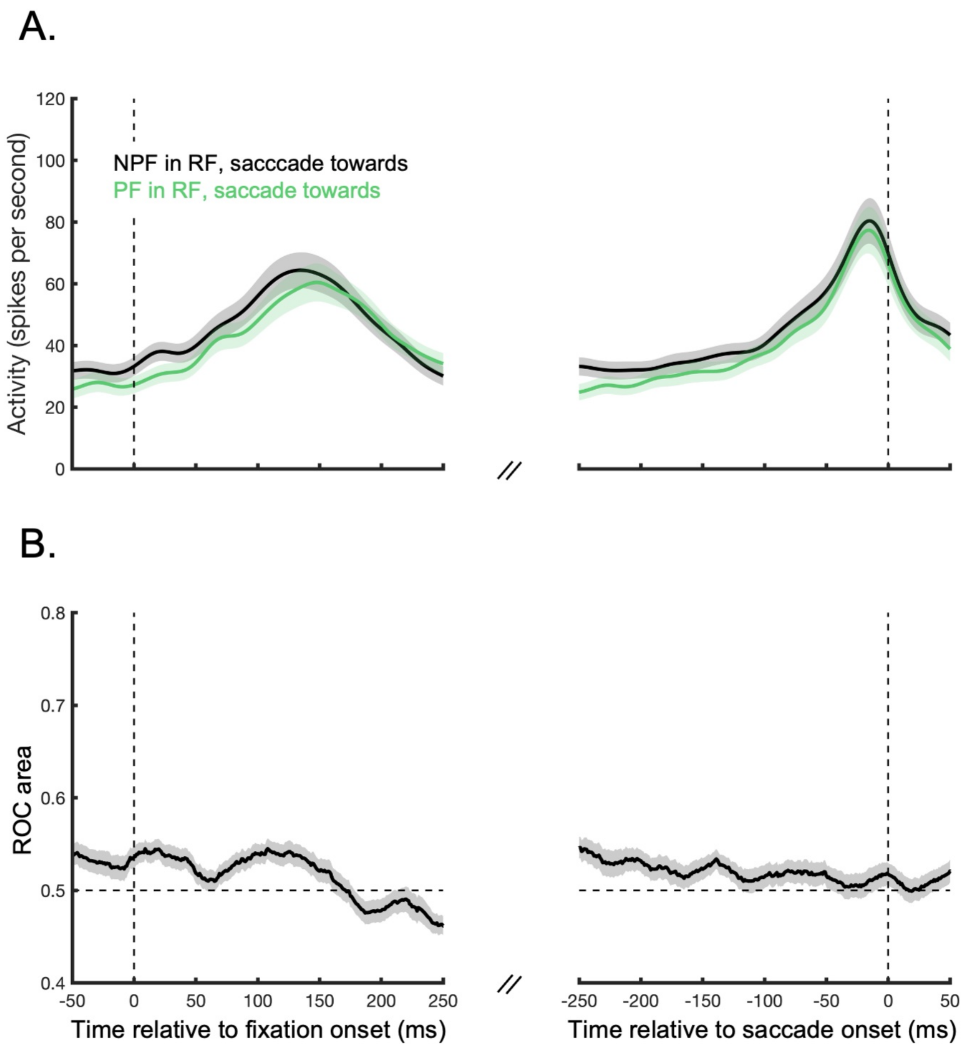


***FIG. S1: A.*** *Activity during NPF-in-RF, saccade towards (black) and PF-in-RF, saccade-towards (green) fixations aligned on fixation onset (left column) and saccade onset (right column). Lines show the mean activity across the 65 neurons analyzed in the main article and shaded regions show +/- one standard error of the mean.* ***B.*** *Functions showing ROC areas as a function of time aligned on fixation onset (left column) and saccade onset (right column) for an ROC analysis that compared NPF-in-RF to PF-in-RF, saccade-towards activity. ROC area functions were obtained for each of the 65 neurons and the lines here show the mean ROC area function across these neurons. The shaded regions shows +/- 1 standard error of the mean. The details of the ROC analysis were the same as the ROC analyses reported in the main article. ROC areas above 0.05 indicate greater NPF-in-RF than PF-in-RF activity and thus suggest inhibitory tagging.*

***Relationship between the improvements in SC target-selection metrics and behavioral search efficiency***

To determine if the improvements in SC target-selection metrics following inhibitory tagging were related to the monkeys' search efficiency, we compared a measure of behavioral search efficiency during each SC recording session to the improvement in selection metrics for the SC neuron recorded during that session. More specifically, for our measure of search efficiency, we computed the mean fixation duration preceding saccades to NPF discs. Shorter fixation durations indicated greater search efficiency because (all else being equal) they enabled the fixation of a greater number of discs and thus increased the probability of reward. This measure was computed on the basis of all saccades to NPF discs for each SC recording session and then compared to the difference in discrimination times and magnitudes yielded by the NPF-re-PF and NPF-re-NPF ROC analyses for the SC neuron(s) recorded during that session. For this analysis, we only considered recording sessions in which the monkey received the reward on at least 20% of all trials, to ensure that we only considered behavioral data from sessions in which the monkey was motivated and on task. Moreover, as in the analyses reported in the main article, we only considered SC neurons that discriminated selected from unselected RF discs for both the NPF-re-NPF and NPF-re-PF analyses, because our interest was the shift in selection metrics across these two analyses.

Figure S2 shows the results of the analysis. Each panel shows the relationship between the mean fixation duration preceding saccades to NPF discs for a particular recording session and the difference in a particular selection metric (discrimination time or magnitude) for the SC neuron(s) recorded during that session. Figure S2A-B show the difference in selection metrics for the fixation-onset-aligned ROC analyses, whereas Fig. S2C-D show the difference in selection metrics for the saccade-onset-aligned ROC analyses. As can be seen, there was a significant relationship between the mean fixation duration preceding saccades to NPF discs and the difference in selection metrics for SC neurons in all cases, suggesting that improvements in SC target selection following inhibitory tagging contributed to efficient behavior. Consider, for example, Fig. S2A. What this plot shows is that shorter fixation durations preceding saccades to NPF discs were related to larger decreases in discrimination times following inhibitory tagging ($R^2$ = .098, $P$ = .029). Similarly, Fig. S2B shows that shorter fixation durations preceding saccades to NPF discs were related to larger increases in discrimination magnitudes following inhibitory tagging ($R^2$ = .177, $P$ = .003).

Figure S2C-D show the same two trends but for the saccade-onset-aligned selection metrics (for the relationship between fixation durations and difference in discrimination times: $R^2$ = .311, $P$ < .001, Fig. S2C; for the relationship between fixation durations and difference in discrimination magnitudes: $R^2$ = .114, $P$ < .024, Fig. S2D).

In summary, the magnitudes of the measured shifts in discrimination time and discrimination magnitude were correlated with shifts in saccade latency to NPF discs, and thus we conclude that the target-selection benefits observed at the level of the SC likely contributed to efficient search behavior in the search task.

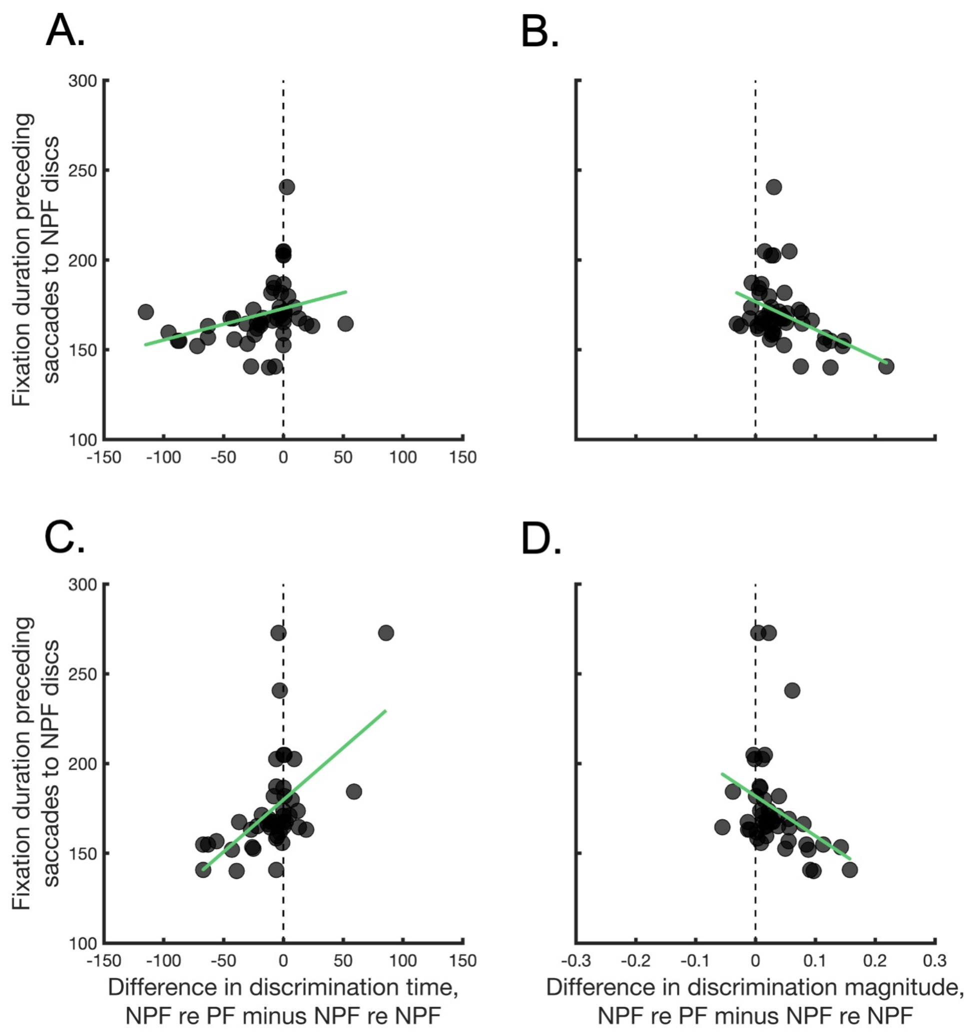


***FIG. S2:*** *Relationship between a measure of search efficiency (i.e., the mean fixation duration preceding saccades to NPF discs) within individual SC recording sessions and the improvements in target-selection metrics for individual SC neurons recorded during those sessions.* ***A.*** *Relationship between mean fixation durations preceding saccades to NPF discs and the difference between NPF-re-PF and NPF-re-NPF fixation-onset-aligned discrimination times (i.e., NPF-re-PF discrimination time minus NPF-re-NPF discrimination time). Negative values on the x-axis represent improvements in the speed of SC target selection following inhibitory tagging.* ***B.*** *Relationship between mean fixation durations preceding saccades to NPF discs and the difference between NPF-re-PF and NPF-re-NPF fixation-onset-aligned discrimination magnitudes (i.e., NPF-re-PF discrimination magnitude minus NPF-re-NPF discrimination magnitude). Positive values on the x-axis represent improvements in the strength of selection signals following inhibitory tagging.* ***C-D.*** *As in A-B but for the saccade-onset-aligned selection metrics. The green line in each panel shows a linear least-squares fit to the data.*
